# Analysis of higher order interactions quantifies co-ordination in the epigenome and reveals novel biological relationships in Kabuki syndrome

**DOI:** 10.1101/2024.03.11.584387

**Authors:** Sara Cuvertino, Terence Garner, Evgenii Martirosian, Bridgious Walusimbi, Susan J. Kimber, Siddharth Banka, Adam Stevens

## Abstract

Complex direct and indirect relationships between multiple variables are a characteristic of all natural systems and are defined as higher order interactions (HOIs). Traditional differential and network analyses fail to account for the richness of omic datasets and miss HOIs. We investigated genome-wide peripheral blood DNA methylation data from Kabuki syndrome type 1 (KS1) and control individuals, identified 2,002 differentially methylated points (DMPs), and inferred 17 differentially methylated regions, which represent only 189 DMPs. We followed these results with quantification of HOIs by applying hypergraph network models on all the CpGs in the two datasets and revealed differences in co-ordination of the DMPs along with lower entropy and higher co-ordination of the peripheral epigenome in KS1 implying reduced network complexity. We demonstrate that the hypergraph approach captures substantially more information, enables factoring trans-relationships, and identifies biologically relevant pathways that escape the standard analyses. These findings construct the basis of a suitable model that is not computationally intensive for the analysis of the organisation of the epigenome in rare diseases. This approach can be applied to other types of omic datasets, and to other fields of science and medicine to investigate mechanism in big data.

## INTRODUCTION

Omic datasets, such as epigenomics and transcriptomics, are usually interpreted using a differential analysis approach, which treats each variable independently. A complementary approach is a network-based analysis, which models biological systems as pairs of interacting variables^1,2^. Network-based models can identify core clusters of genes that are likely to be mechanistically relevant but can miss complex direct and indirect relationships^3^ especially those between multiple variables^4^.

Complex systems can rarely be captured by their pairwise dynamics alone^5,6^. Rather, natural systems demonstrate relationships between multiple variables^7^ known as higher order interactions (HOIs). HOIs can be measured by hypergraphs, which are a generalisation of a graph (network) in which an edge can join any number of vertices. Hypergraphs can reveal mechanistic insights that can escape traditional analyses^8–10^, including the impact of both HOIs^5,11^ and direct and indirect interactions. Importantly, hypergraph models are viewed as mechanistic and do not rely on qualitative assessment of gene ontology to establish function^8–10,12^.

Elements of complex systems are co-ordinated by higher order interactions^9^. This coordination can be assessed using entropy^13^, which measures network structure, combined with measuring network path directness. They have been used to examine early human embryology^14,15^ and hypergraph topology has been used to assess mechanism in neural, ecological and social systems^11^. Recently, efficient imputation of multi-tissue and cell-type gene expression has been achieved using a hypergraph approach^16^. However, hypergraphs have not yet been used to assess mechanistic relationships in human diseases. In this study, we have tested the hypergraph approach in context of epigenomic data from a rare disease, Kabuki syndrome (KS) type 1.

KS is one of the commonest Mendelian histone lysine methylation disorders^17^. Most KS cases are caused by heterozygous loss-of-function variants in H3K4 methyltransferase 2 D (*KMT2D)* (KS1, OMIM#147920), while less than 5% of cases are caused by X-linked *KDM6A* variants (KS2, OMIM#300867)^18–20^. By regulating enhancer and promoter elements, KMT2D controls the access of transcription factors and other proteins to DNA and therefore controls gene transcription and regulation^21–25^. Clinically, KS is characterised by distinct facial dysmorphism, intellectual disability, developmental delay, and a range of internal organ malformations such as congenital heart defects, skeletal defects, cleft palate and genitourinary malformations^26^. In addition, affected individuals are susceptible to several functional anomalies (e.g., endocrine disorders, deafness) and immune defects^26–28^.

DNA methylation can act in both a *cis-* and *trans-* manner in the control of gene expression^29^, however even those studies which investigate trans-interactions do not consider higher order effects. Here, we have integrated KS1 and control genome-wide DNA methylation data from three studies to identify novel differentially methylated CpGs (differentially methylated points [DMPs]) and differentially methylated regions (DMRs). To identify higher order interactions in the KS1 epigenome we have generated hypergraph models of observed methylation patterns, assessed epigenomic co-ordination by quantifying the entropy of those network models, and investigated the directness of the relationships between epigenomic points defined by them (**Figure 1**). Our approach reveals multiple novel, biologically relevant and likely causal relationships in KS1. Overall, we show that hypergraphs allow investigation of the entire epigenome, including co-ordination over genomic distance, in the context of a rare disease, and provides a new quantifiable framework to reveal causal relationships.

**Figure 1.**
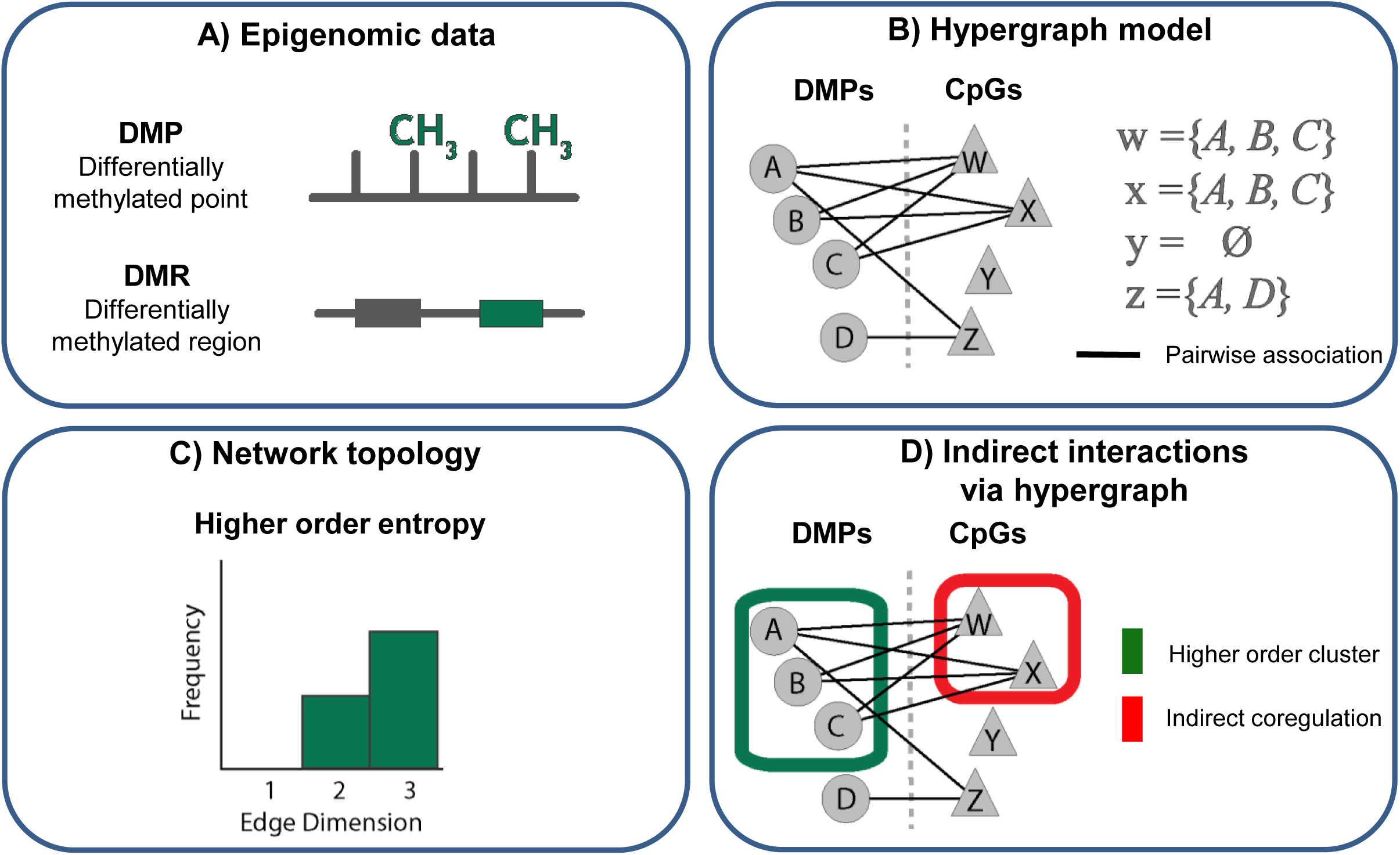
Experimental design to assess the co-ordination of peripheral blood DNA methylation associated with Kabuki syndrome. The core features of the experimental design used to assess differences in the co-ordination of the epigenome between KS1 and controls. **A)** Peripheral blood DNA methylation is used to identify differences in the epigenome between KS1 and controls. Differentially methylated points (DMPs) and regions (DMRs) are defined using a statistical approach. **B)** Pairwise associations are defined between DMPs (*A-D*) and all remaining CpGs (*W-Z*). These bipartite network models distinguish HOIs from pairwise associations in the epigenome. Associations with CpGs which are common between DMPs can be considered higher order interactions between DMPs, or hyperedges, described here by sets *w-z*. **C)** To measure co-ordination of these higher order networks, an indicator of function, Shannon entropy is calculated on the distribution of edge dimensions. **D)** Hypergraph models are generated from these bipartite structures and refine clusters of co-ordinated DMPs (green box) as well as implicating a wider set of CpGs as potentially indirectly co-regulated (red box).

## RESULTS

### DMR analysis using integrated dataset identifies new biological processes disrupted in KS1

Firstly, we set out to identify the DMRs in KS1 compared to controls. We identified 417,217 common probes across the three peripheral blood DNA methylation array datasets to enable comparison between data from 22 individuals with KS1 and 138 controls (**Figure 2A**). PCA analysis performed using only DMPs demonstrated segregation of KS1 individuals from controls suggesting robustness of our data integration (**Figure 2B**).

**Figure 2.**
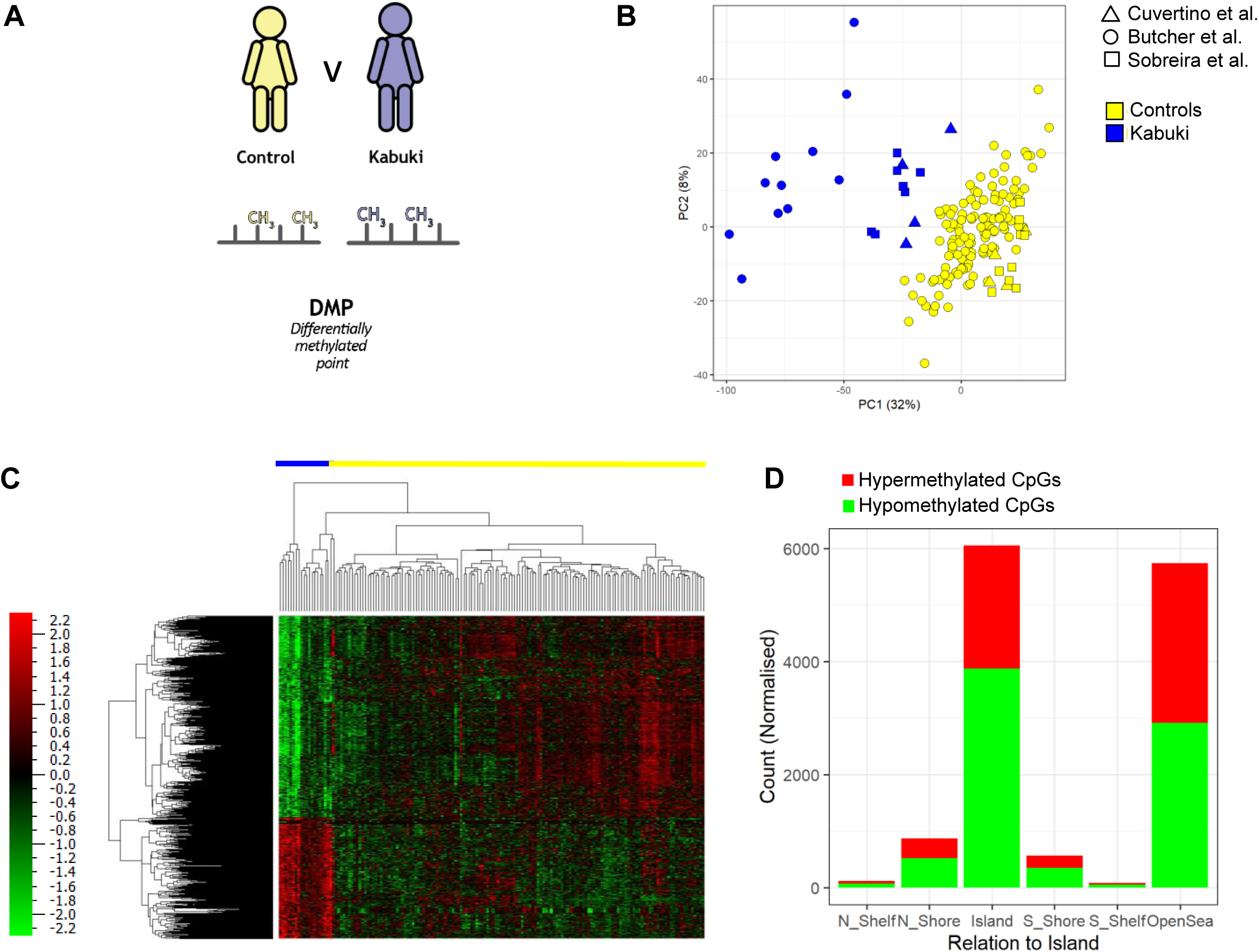
Peripheral blood DNA methylation reveals differences between KS1 and controls. **A)** Differentially methylated points are identified between KS1 and control patients. Control samples are indicated in yellow, KS1 in blue. **B)** Principal Component Analysis (PCA) of the peripheral blood DNA methylome demonstrates variation between KS1 and control samples. Shapes are used to differentiate the studies from which data were drawn: circles represent samples from *Butcher et al.*, squares represent *Sobreira et al.* and triangles *Cuvertino et al*. **C)** A heatmap of relative methylation of DMPs (FDR<0.0001). Hierarchical clustering of samples and DMPs is represented by dendrograms on the columns and rows respectively. Nature of samples is indicated by the coloured bar above the column dendrogram. Relative hypo- and hypermethylation are represented by green and red in the heatmap, respectively. **D)** A bar graph demonstrates the number of DMPs that are hypo- and hypermethylated in KS1 compared to controls (FDR<0.0001). DMPs are grouped by proximity in base pairs to CpG islands.

From this dataset, we identified 2,002 DMPs in the KS1 samples compared to the controls (FDR<1x10^-4^). Of these, in KS1 753 DMPs were hypermethylated and 1,249 were hypomethylated (**Figure 2C**). Our analysis identified more DMPs compared to previous studies^30,31^. DMPs were found to be located predominantly in CpG islands and open sea (4 Kb from CpG islands) regions (**Figure 2D**).

Using these DMPs, we identified 208 DMRs (> 7 annotated DMPs) between KS1 and controls (87 hypermethylated and 121 hypomethylated, FDR<1x10^-3^) (**Figure 3A**). Of these, 17 DMRs (11 hypermethylated and 6 hypomethylated) passed the cut-off of FWER<0.05 (**Table 1**) (**Figure 3B**). Our analysis validated 3 previously identified DMRs (*HOXA4, HOXA5, HOXA6*)^30,31^ for KS1, and identified 14 additional DMRs, even though the cut-offs used here were more stringent than the previous studies^30,31^.

**Figure 3.**
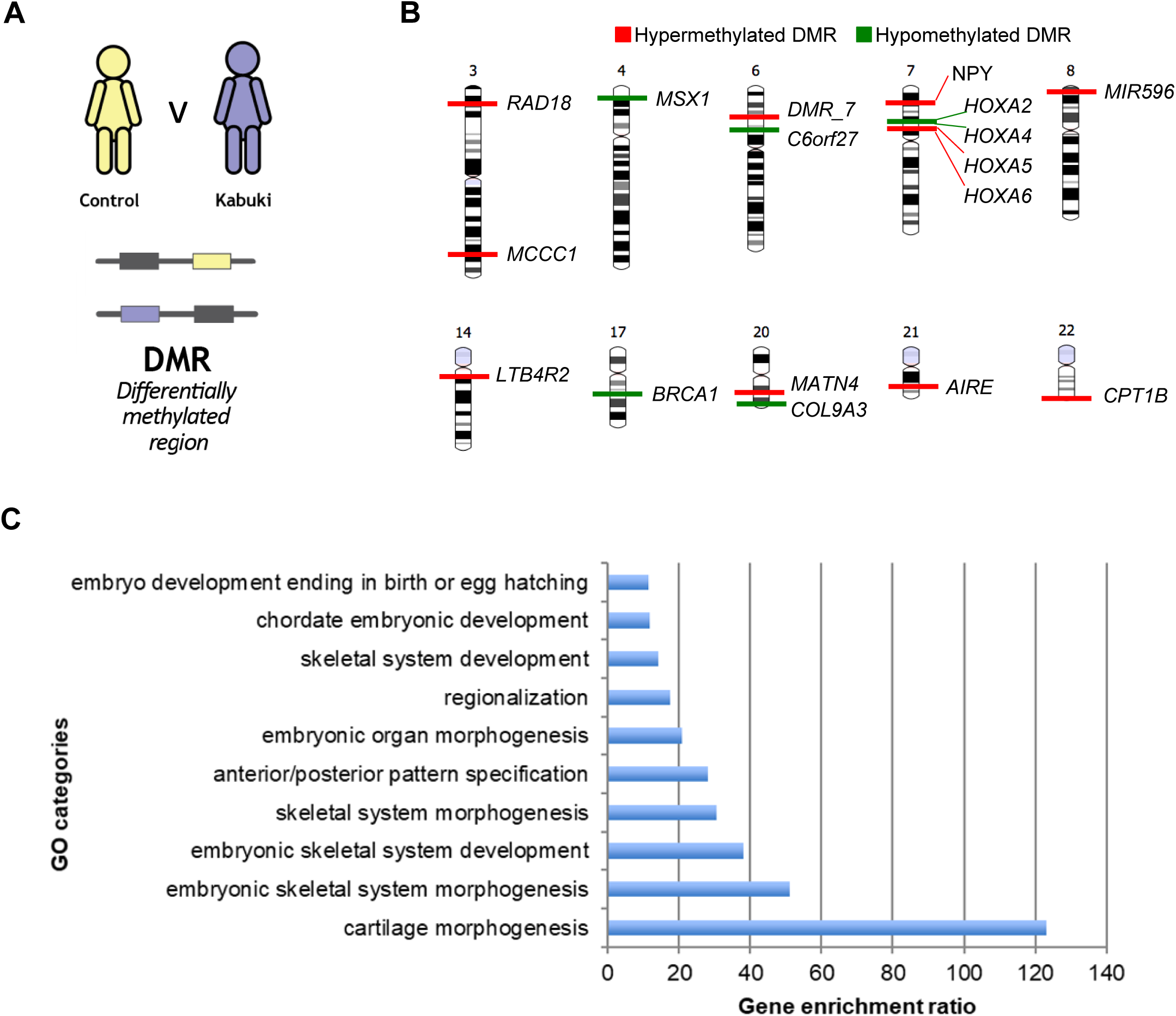
KS1 results in differences in methylated regions between KS1 and controls. **A)** Differentially methylated regions are identified between KS1 and control patients. Control samples are indicated in yellow, KS1 in blue. **B)** Ideograms show the 17 DMRs (FWER<0.05) identified using ChAMP analysis. Hypomethylated DMRs (11) are shown in green and hypermethylated DMRs (6) are in red. **C)** List of gene ontology categories performed on the 17 significant DMRs (11 hypermethylated and 6 hypomethylated) which passed the cut-off of FWER<0.05 (WebGestalt; FDR<0.05).

**Table 1.** Significant DMRs identified in KS1. List of DMRs including chromosome (Chr), gene annotation, relative methylation value, P-value and family wise error rate (FWER) corrected P-value. DMRs with FWER<0.05 are included.

| DMR | Chr | Genes | Value | P-value | FWER |
| --- | --- | --- | --- | --- | --- |
| DMR_1 | chr7 | HOXA5 | 7.87 | <2.2x10<sup>-16</sup> | <2.2x10<sup>-16</sup> |
| DMR_2 | chr14 | LTB4R2 | 8.76 | <2.2x10<sup>-16</sup> | <2.2x10<sup>-16</sup> |
| DMR_3 | chr8 | MIR596 | 9.88 | <2.2x10<sup>-16</sup> | <2.2x10<sup>-16</sup> |
| DMR_4 | chr7 | HOXA4 | -6.65 | <2.2x10<sup>-16</sup> | <2.2x10<sup>-16</sup> |
| DMR_5 | chr7 | HOXA6 | 8.16 | <2.2x10<sup>-16</sup> | <2.2x10<sup>-16</sup> |
| DMR_6 | chr7 | HOXA2 | -6.26 | <2.2x10<sup>-16</sup> | <2.2x10<sup>-16</sup> |
| DMR_7 | chr6 | - | 6.36 | 6.86E-06 | 0.001 |
| DMR_8 | chr7 | NPY | 7.55 | 4.80E-05 | 0.001 |
| DMR_9 | chr4 | MSX1 | -6.64 | 5.72E-05 | 0.001 |
| DMR_10 | chr20 | MATN4 | 6.33 | 8.23E-05 | 0.002 |
| DMR_11 | chr17 | BRCA1 | -6.01 | 8.23E-05 | 0.005 |
| DMR_12 | chr22 | CPT1B | 5.98 | 1.23E-04 | 0.008 |
| DMR_13 | chr6 | C6orf27 | -5.86 | 8.46E-05 | 0.012 |
| DMR_14 | chr21 | AIRE | 6.18 | 1.67E-04 | 0.02 |
| DMR_15 | chr3 | RAD18 | 84.13 | 9.60E-05 | 0.041 |
| DMR_16 | chr20 | COL9A3 | -4.47 | 1.03E-04 | 0.045 |
| DMR_17 | chr3 | MCCC1 | 9.30 | 1.23E-04 | 0.05 |

We then performed gene ontology analysis using the 17 significant DMRs. This showed genes within these regions to be associated with embryonic skeletal system morphogenesis and development (*HOXA2, HOXA4, HOXA5, HOXA6, MSX1, MATN4, COL9A3*), anterior/posterior pattern specification and embryonic organ morphogenesis (*HOXA5, HOXA4, HOXA6, HOXA2, MSX1*) (**Figure 3C**).

Next, using Encode data we analysed the epigenetic landscape of the DMRs in human blood cells (B cells, CD133 haematopoietic stem cells, and Neutrophils) using the UCSC genome browser (**Supplementary Figure S1 and Supplementary table 1**). These cells were chosen because our DNA methylation dataset is from peripheral blood DNA samples. Seven out of 17 DMRs showed enrichment for H3K4me3 and H3K27Ac indicating transcriptional activity in these cells. As DNA methylation regulates gene expression, we also evaluated the level of expression of the genes corresponding to the identified DMRs in RNAseq datasets (GSE126167, GSE126166)^32^ from human or mouse KS1 cell models. In murine HT22 cells with homozygous deletion of the catalytic SET domain of *Kmt2d*, *Hoxa2* and *Aire* were differentially expressed while *Npy* and *Col9a3* were down-regulated in EdU+ DG nuclei of *Kmt2d*^+/-βgeo^ mice compared to controls^32^.

Collectively, these results reveal evidence of disruption of several disease relevant biological processes in peripheral blood derived epigenome of KS1, and thus demonstrate the power of data integration in revealing novel mechanistic insights.

### Hypergraph analysis identifies differences between biological pathways enriched in KS1 and control DMPs

Although informative, the 17 significant DMRs collectively represent only 189/2,002 significant DMPs. Hence, the DMR analysis ignored the functional relevance of >90% (1,817 out of 2002) of the significant DMPs. This is especially true for DMPs present in open sea regions, where CpGs and probe density are lower, thus reducing the chances of identifying DMRs (**Figure 2D**). We, therefore, wanted to interrogate the mechanistic relevance of all significant DMPs. However, methylation status of multiple DMPs or CpGs could affect the methylation status of multiple DMPs or CpGs. In other words, DMPs are likely to have a complex co-ordination with other DMPs and other CpGs. Hence, treating DMPs as independent variables and simply looking for enrichment of genes in the DMP datasets could be misleading. We, therefore, employed a hypergraph approach to identify groups of DMPs exhibiting HOI utilizing the entire epigenomic dataset^9,10,12^. In this context, the hypergraph summarizes correlations between each DMP and all remaining CpGs to infer relationships between DMPs; larger numbers of shared correlations between DMPs indicate more likely higher order (and therefore causal) relationships.

Applying this hypergraph approach showed that 986/2,002 DMPs in KS1 and 1,036/2,002 DMPs in controls were highly co-ordinated (appearing as heatmap clusters in **Figure 4A and 4B**). This means that methylation of these DMPs in KS1, or in controls, is highly interrelated. These results were validated using an independent approach of non-negative matrix factorisation (**Supplementary Figure S2**). Interestingly, 822/2,002 highly co-ordinated DMPs were found to be shared across both KS1 and control samples. This implies that 164/986 KS1-exclusive DMPs represent likely new disorder-specific interactions and 214/1,036 controls DMPs represent the interactions that are likely lost in KS1 (**Figure 4C**).

**Figure 4.**
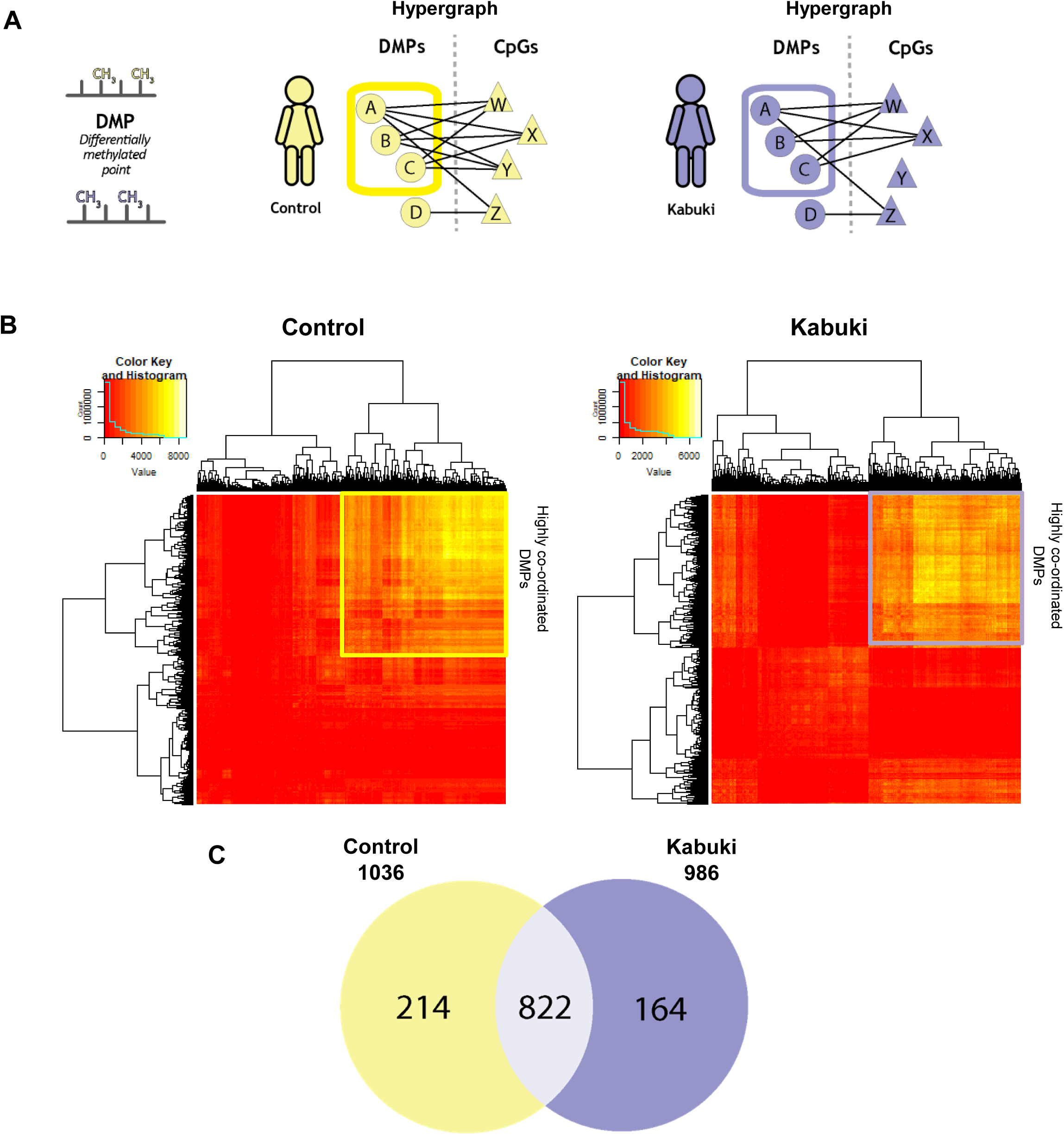
Clustering hypergraph adjacency matrices reveals higher order interactions between DMPs which distinguish KS1 and controls. **A)** Pairwise relationships between DMPs and CpGs can be summarised as higher order interactions between pairs of DMPs using a hypergraph approach. **B)** Heatmaps of hypergraph adjacency matrices for control (L) and KS1 (R). Red to yellow colouring in the heatmap represents increasing dimensionality of the hyperedges between pairs of DMPs. Hierarchical clustering of the hyperedges reveals a central cluster of highly co-ordinated DMPs in controls (yellow box) and KS1 (purple box) associated to one another by higher order interactions. **C)** Venn diagram demonstrating the overlap of DMPs in the central cluster of the hypergraphs.

Next, we performed gene ontology analysis on the 986 KS1 co-ordinated DMPs and identified an enrichment for genes (P<0.05, FDR>0.05) associated with connective tissue development (*TGFB1*), cartilage development (*TGFB1*) and neuronal migration (*TGFB1, NAV1*). In contrast, gene ontology analysis performed on the 1,036 control co-ordinated DMPs demonstrated an association with response to interleukin-17 and vitamin D (**Supplementary Figure S3**).

Collectively, these results demonstrate the utility of hypergraph analysis to identify biologically relevant pathways from DNA methylation data that can escape the standard DMR-based analysis.

### Shannon entropy analysis shows differences in the network topologies of KS1 and control DMPs

The hypergraph analysis showed that a large proportion of the clusters highly co-ordinated DMPs (822 DMPs) were shared between KS1 and controls, implying that although these CpGs are significantly differentially methylated in KS1, they remained highly co-ordinated in both KS1 and controls. Although the DMPs may remain highly co-ordinated in both sets, the nature (i.e. the direction or the strength) of these co-ordinations may be different in KS1 and controls. These differences are represented in the hypergraph model as the ‘network topology’. Differences in network topology can be measured using Shannon entropy^33^, which examines the distribution of the number of shared correlations between any pair of DMPs (hyperedge dimensionality) in the hypergraph model (**Figure 5A**). We, therefore, performed an iterative analysis of comparing Shannon entropy within the central cluster of the adjacency matrix and randomly selected CpGs of hypergraphs with 1,000 randomly generated simulations of 100 DMPs. We observed a lower entropy in the KS1 epigenome compared to controls, implying more ordered DMP co-ordination in the disorder (**Figure 5B**).

**Figure 5.**
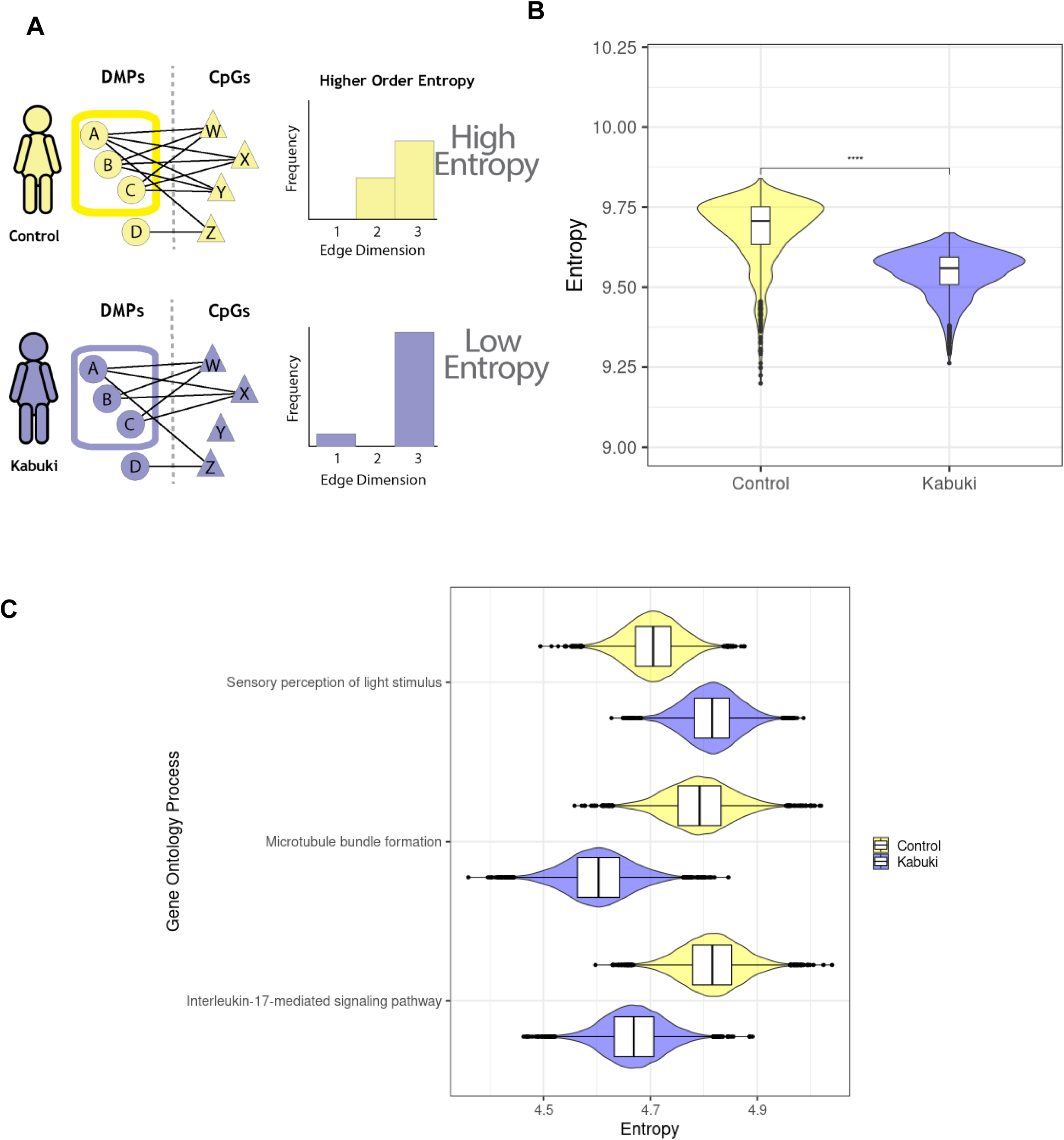
Topology of control and KS1 hypergraphs highlight differences in co-ordination. **A)** Co-ordination of the central cluster of the hypergraph in control (yellow) and KS1 (purple) can be quantified using Shannon entropy. Edge dimension is the number of shared correlations between a pair of vertices in the hypergraph model; entropy quantifies network structure from the distribution of edge dimensionality, such that a highly uniform network would have low entropy. **B)** Shannon entropy of the hypergraph central cluster of control and KS1 methylome. Presented data are compared to data from 1000 matched iterations. **C)** Difference in entropy for pathways identified as enriched in the central clusters of either the control or KS1 hypergraphs. Bayesian Markov Chain Monte Carlo sampling was performed to enable comparison between control and KS1. Only significantly different pathways (those where the 89% Credible Interval of the difference does not contain 0) are presented.

Next, we assessed the contribution of the pathways enriched in the hypergraph central clusters (**Supplementary Figure S3**) to the overall epigenome entropy using a Bayesian approach (**Supplementary Figure S4**). This analysis shows that three pathways which were differentially enriched in the hypergraph central clusters have significantly different network entropy in KS1 and controls – “sensory perception of light stimulus” (higher entropy in KS1) and “microtubule bundle formation” and “Interleukin-17-mediated signalling” (lower entropy in KS1) (**Figure 5C**). Importantly, none of these pathways were identified using the traditional approach of one-to-one comparison of DMPs or DMRs.

This analysis shows that the KS1 co-ordinated epigenome has a different network topology compared to controls suggesting a more ordered and less diverse co-ordination by the DMPs in KS1.

### Indirect association of co-ordinated DMPs is distinct in KS1 compared to controls in the hypergraph models

The results presented in this manuscript so far are based on analysis of DMPs. However, there may be disorder relevant to differences in the co-ordination of regions of the epigenome even if individual points may not be significantly differentially methylated. The hyperedges^9^ in these models represent all co-ordinated CpGs within the entire epigenome, including those that are peripheral and not just the DMPs (i.e. the set of CpGs being co-ordinated by higher order action), thus capturing indirect associations in the epigenome (**Figure 6A**). We, therefore, quantified the peripheral associations for KS1 and for controls. We detected more peripheral associations in KS1 compared to controls (2,170 and 1,381 CpGs respectively) with minimal overlap between the identified genes (**Figure 6B**).

**Figure 6.**
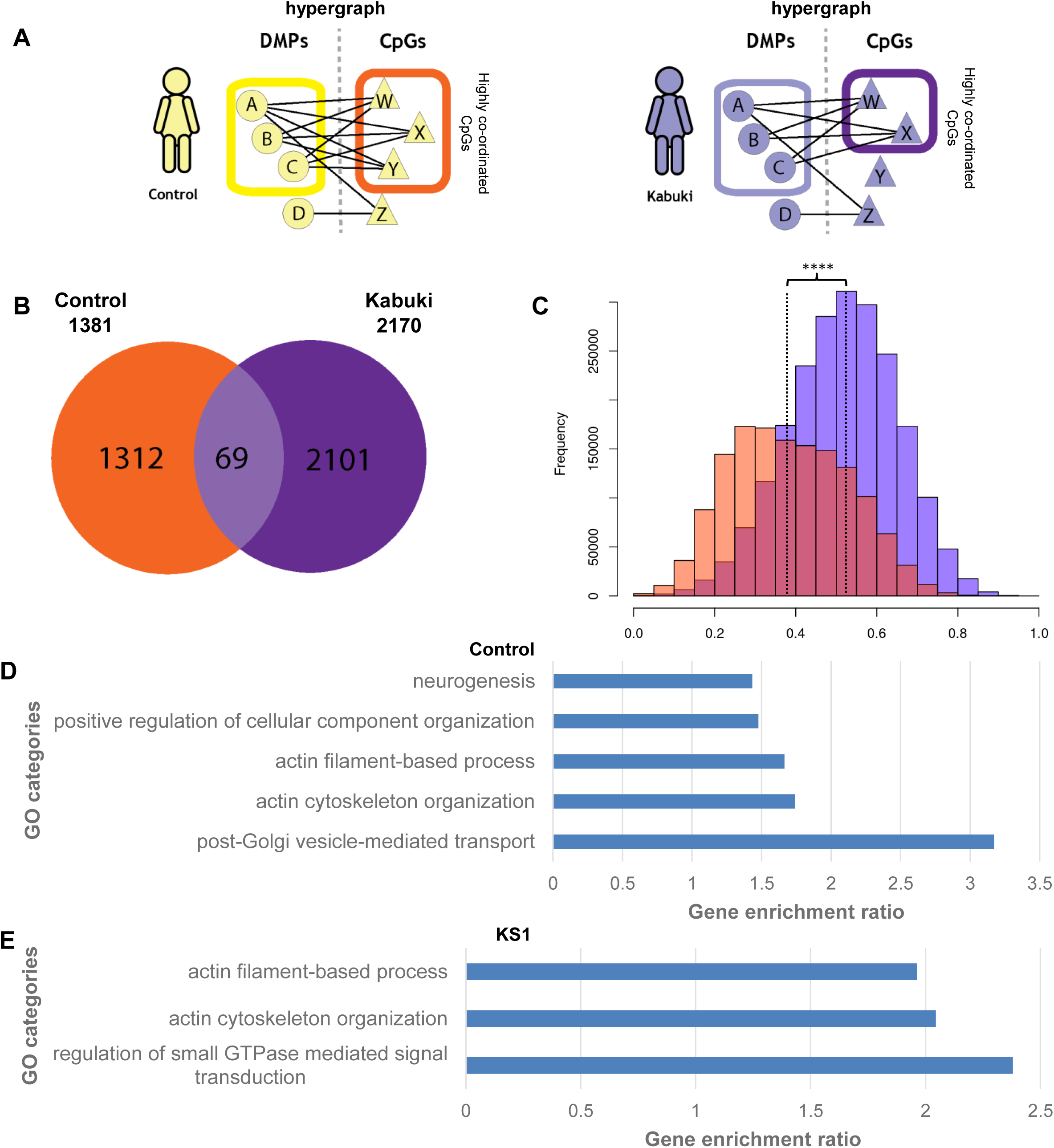
Co-ordinated DMPs demonstrate potential indirect co-regulation of a wider set of CpGs. **A)** In addition to refining clusters of co-ordinated DMPs (yellow/purple box) the hypergraph approach also implicates a wider set of CpGs (orange/dark purple box) as potentially indirectly co-regulated with the co-ordinated DMPs (yellow/purple box). **B)** Venn diagram of the indirectly co-regulated CpGs to demonstrate overlap between control and KS1. **C)** Distribution of correlation values (|r|) between DMPs and indirectly regulated CpGs in control (orange) and KS1 (dark purple). **D-E)** Ontology of nearest gene to indirectly regulated CpGs which were unique to control (D) or KS1 (E) (WebGestalt, FDR<0.01).

To quantify the indirect associations, we calculated the frequency of correlation values (|*r*|) in our CpGs. Comparing these data, we observed that the correlation values in KS1 were 1.3-fold higher compared to controls (P<1x10^-9^) (**Figure 6C**). Peripherally related CpGs in KS1 were enriched for gene ontologies related to small GTPase signal transduction (FDR<0.01) (**Figure 6D**) indicating that these indirect associations are likely gained in KS1. Peripherally related CpGs in controls were enriched for vesicle mediated transport and neurogenesis (FDR<0.01) (**Figure 6E**) indicating that these indirect associations are likely lost in KS1. Gene ontologies related to actin cytoskeleton organization were enriched in both KS1 and controls. These results highlighted a greater variety of indirect associations of CpGs in KS1 compared to controls, implying major functional shifts in the wider epigenome between KS1 and controls.

## DISCUSSION

We have analysed the impact of higher order interactions within the epigenome in a rare disease context to aid the understanding of disease mechanism (summarised in **Figure 7**). Using the DMR based analysis, DMPs-based hypergraph, entropy studies, and entire epigenomic based indirect co-ordination analysis we identified several disease relevant genes, regions and pathways with mechanistic links.

**Figure 7.**
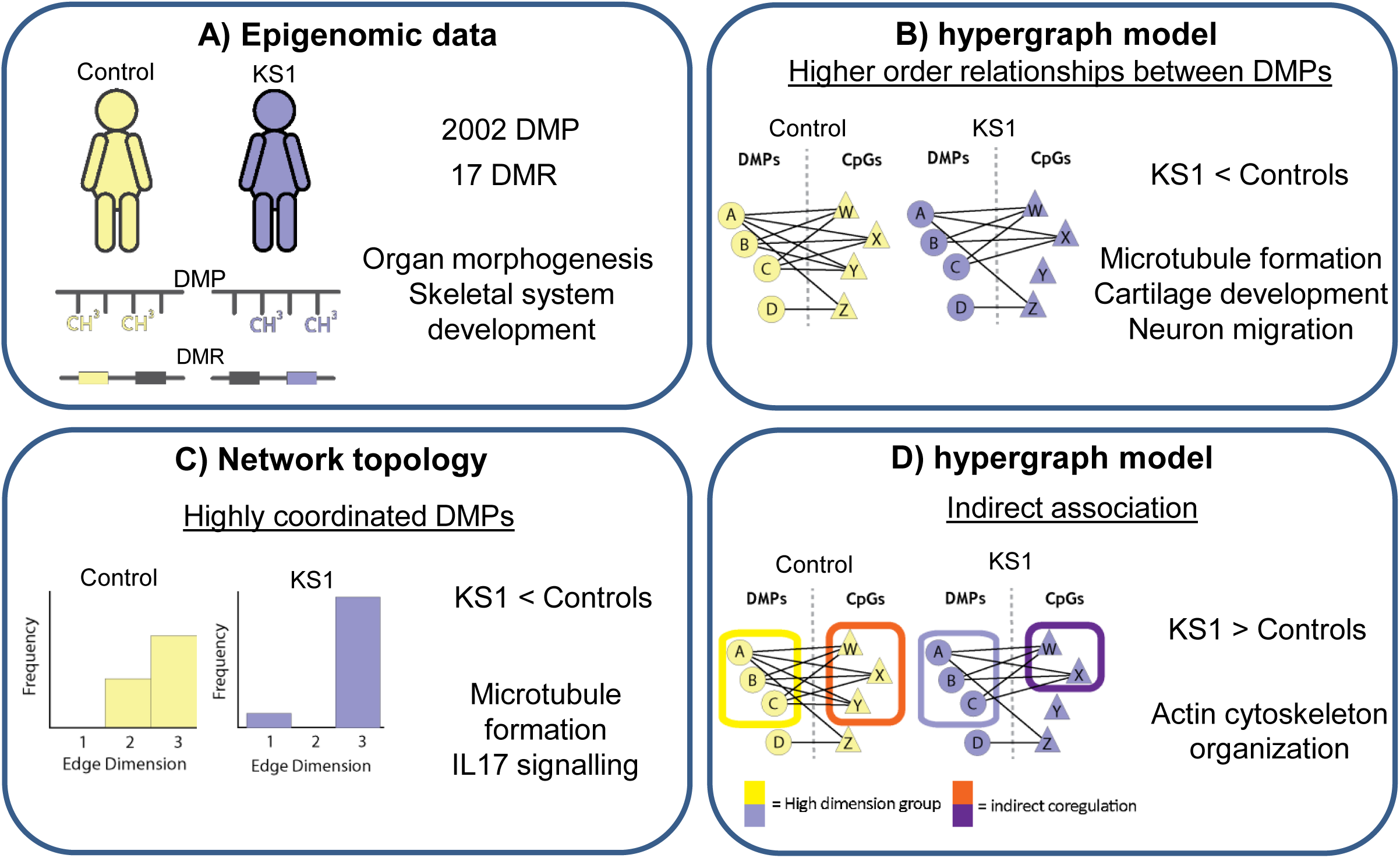
Analysis of blood DNA methylation reveals single CpG and regional differences, as well as differences in the co-ordination of the epigenome between KS1 and control. **A)** Statistical analysis highlights differences between KS1 and control in methylation of genes associated with morphology. **B)** HOIs between DMPs reveal a cluster of co-ordinated DMPs, present in both KS1 and control, in genes associated with organ development. **C)** Quantification of the co-ordination of DMPs demonstrated a lower entropy and therefore more defined co-ordination between DMPs in KS1 than control. **D)** The co-ordinated DMPs were indirectly associated with a wider range of CpGs in KS1 than controls. Differences in co-ordinated DMPs and indirectly implicated CpGs were associated with development and cellular organization.

Four DMRs correspond to *HOXA* genes, which play a fundamental role in embryonic development and in the anterior to posterior pattern specification^34^ (**Figure 3**). Two HOXA DMRs, corresponding to *HOXA4* and *HOXA2,* were found to be hypomethylated and two corresponding to *HOXA5* and *HOXA6* were hypermethylated. HOXA2 is involved in the dorsal-ventral patterns of neuronal development in the rostral hindbrain^35^ and *HOXA2* variants cause microtia, hearing impairment and cleft palate (OMIM 612290)^36,37^. *HOXA4 is* involved in the anterior transformations of the dorsal aspects of components of the vertebral column^38^. HOXA5 is important in the patterning of the cervico-thoracic region and regulates organ

development such as that of fore limb specification, cartilage, lung and gut^39^. HOXA6 is also important for hematopoietic cell proliferation and self-renewal^40^. We detected hypermethylation in the DMR corresponding to *MSX1* (**Figure 3**), which has roles in limb-pattern formation and craniofacial development, particularly odontogenesis^41^. *MSX1* variants cause Witkop type ectodermal dysplasia 3 (OMIM 189500), orofacial cleft type 5 (OMIM 608874) and selective tooth agenesis (OMIM 106600)^42,43^. Dysregulation of these genes might explain the broad range of organ malformations and anomalies such as congenital cardiac defects, renal malformations and dental anomalies observed in KS1^26^.

We observed hypomethylation of DMR representing *NPY* (**Figure 3**), which encodes a neuropeptide that is widely expressed in the central nervous system and influences many physiological processes, including cortical excitability, food intake, and cardiovascular function^44,45^. Gene ontology analysis of co-ordinated DMPs in KS1 identified DMPs representing *TGFB1, NAV1* (**Figure 3**) that are involved in neuronal migration. We detected loss of co-ordination in peripherally related CpGs to be enriched for vesicle mediated transport and neurogenesis in Kabuki. Thus, dysregulation of these genes and pathways and processes may contribute to the neurodevelopmental and neurological phenotype of KS1 such as intellectual disability, with visuospatial construction, perception, memory, and language impairment^46^.

Hypermethylation of *LTB4R2* (**Figure 3**), a low-affinity receptor for leukotrienes^47^, and hypermethylation of *AIRE,* a transcriptional regulator that interacts with the transcriptional coactivator CREB protein^48^ were observed. These genes will be good targets to interrogate the mechanism of immune phenotypes in patients with KS1. We also detected loss of co-ordination and lower entropy of DMPs representing interleukin-17 in KS1. Dysregulation in these genes and pathways may contribute to the immune phenotypes of KS1 such as increase infection susceptibility and hypo-gammaglobulinemia^49^.

We identified DMRs corresponding to *COL9A3* and *MATN4* (**Figure 3**), which were hypo and hypermethylated, respectively. COL9A3 is a structural component of hyaline cartilage and the vitreous of the eye^50^. MATN4 is part of the extracellular matrix of various tissues in particular cartilage^51^. *COL9A3* variants cause Stickler syndrome type VI (OMIM 620022) characterised by hearing loss and skeletal abnormalities^52,53^. Dysregulation in these genes and pathways may contribute to the skeletal defects and joint hypermobility that are frequent in KS1.

The mechanism of generation of the abnormal DNA methylation pattern in individuals with KS1 remains to be examined. It could be a direct consequence of altered histone modifications and increased or decreased access to DNA methyltransferases and demethylase enzymes. Of note, KMT2D marks H3K4 in active enhancer regions, whereas CpG islands usually correspond to gene promoter regions. Hence, this raises alternative possibilities, such as that DNA methylation changes may be a secondary consequence of alterations in wider chromatin architecture or the 3D genomic architecture. It is also possible that these changes may be a consequence of subtle changes in cellular signalling, leading to alterations in developmental trajectories. It is notable that analysis based on DMRs derived from blood cells revealed dysregulation of genes and pathways that may contribute to non-blood phenotypes. Interestingly, we detected the greatest changes in DMRs associate with the GO term cartilage morphogenesis (**Figure 3C**). This may be explained by the shared embryonic mesodermal origin of blood and the skeleton.

The DMPs based hypergraph modelling approach employed here measures system-wide co-ordination of epigenomic features. We identified a subset of DMPs with higher order co-ordination of their methylation levels, indicating co-ordination in their function (**Figure 4**). Interestingly, the hypergraph models of both KS1 and controls contain this same group of co-ordinated DMPs (**Figure 4C**) suggesting similar functional relationships between differentially methylated CpGs in both groups. Despite this similarity, the hypergraph models show a difference in epigenomic network structure between groups, highlighted by a reduction in entropy within KS1 (**Figure 5**). Lower entropy of omic networks is associated with ageing^54,55^ and advancing cellular differentiation^32,33,56^.

Along with these DMPs, however, this approach highlighted a wider set of CpGs-hyperedges (shared correlations between any pair of DMPs) which were important in establishing the relationship between the co-ordinated DMPs. These CpGs were not differentially methylated but were implicated by the hypergraph model as important and still demonstrated relevant biological context. This analysis has highlighted possible disruption in microtubule formation which are essential for cell division, intracellular transport, and cell morphology and organization. This may affect the mechanical properties in the cell and in the nucleus as supported by Negri et al^57^. The importance of these CpGs in the higher order organisation of the epigenome may help to explain differences observed in individuals with genetic variants which are not accounted for by DMPs.

Gene ontology analysis may have limitations in DMP and DMR analysis as this emphasises *cis-* based gene regulation. Hypergraph analysis factors in *trans-* regulatory gene regulation because as it uses the entire epigenome not just that present within genes or near genes. The overlap between hyperedges highlights chains of higher order interactions within the epigenome that may be considered as quantification of co-ordination and therefore a measurement of dysregulation rather than just the identification of directly associated disease related genes and pathways.

In conclusion, our study shows that the comparative analysis of peripheral blood DNA methylation reveals fundamental differences in the apparent functional activity of the epigenome. This study has identified novel candidate genes and pathways for KS1 disease pathology. Importantly, hypergraph approaches have not yet been used to assess mechanistic relationships and pathogenicity in single gene rare diseases. Our findings suggest that these approaches have the potential to reveal additional disease relevant mechanistic insights, which can be quantified, using epigenomes of rare disorders when compared with the traditional one-to-one differential analysis.

## MATERIAL AND METHODS

### Data acquisition

Peripheral blood DNA methylation data from individuals with KS1 and controls was obtained from previously published studies namely Butcher *et al*, Sobreira *et al* and Cuvertino *et al* ^30,31,58^. Of note, we only included data from individuals with variants that could be definitely predicted to cause KS1. Specifically, from the dataset of Butcher *et al*^30^, we excluded 32 individuals with CHARGE syndrome (OMIM#214800)^12^. From the dataset of Sobreira *et al*^31^, we excluded 2 individuals with *KMT2D* variants of uncertain significance and 6 individuals with other variants. From the dataset of Cuvertino *et al*^58^, we excluded 4 individuals with exon 38 and 39 missense *KMT2D* variants that cause a syndrome that is genetically, epigenetically, and phenotypically distinct from 4 KS1 individuals.

### Data processing

All statistical analyses were performed in R 3.4.1 (www.r-project.org)^59^. The studies reported by Butcher *et al* (GSE97362) and Sobreira *et al* (GSE116300) were performed using Illumina Human Methylation 450K (hitherto called 450K arrays)^30,31^ and Cuvertino *et al* used Infinium Methylation EPIC bead chip (Illumina)^58^. In order to make the datasets comparable we removed all non-matching probes between the EPIC and 450K arrays using the Bioconductor package *minfi*^60^. Distribution of cell types within the whole blood samples were estimated and adjusted using the established Houseman method for each individual^61^. Initial data quality was measured by removing the methylated and unmethylated background signal levels exceeding the detection threshold of P>0.01. In addition, to minimize the differences between the samples, we excluded cross-reactive probes^62^, probes on sex chromosomes and those that are age-associated^63–65^. Raw beta values were logit transformed to M values following functional and quantile, within array, normalisations (SWAN)^66^. We used a z score to normalise data for the study.

### Identification of significant DMPs and DMRs in KS1

Methylation data were analysed using the ChAMP^67^ pipeline in R 3.4.1. DMPs were identified using a linear modelling approach to identify differentially methylation levels between groups of samples. DMRs were identified using the Bumphunter^68^ method which defined DMRs as genomic regions with more than 7 DMPs, with a maximum gap of 300bp between DMPs. This technique performs a randomised permutation approach to identify regions with greater than expected number of DMPs. A false discovery rate modified P<1x10^-4^ was used to identify significantly DMPs between KS1 individuals and controls. Z-scores of all methylated positions (3166 probes) were analysed using Qlucore Omics Explorer 3.4 (Qlucore, Lund, Sweden). Principal components analysis (PCA) was used to visualise clustering of the samples based on DMPs between groups. A heatmap was generated using R package heatmap.2 to examine clustering of DMPs using the Euclidean metric. Relation to the island distribution graph was generated in R 3.4.1.

### Statistical analysis

Significant differential methylation was defined as those CpGs with a false-discovery rate modified (FDR) P<1x10^-4^. FDR correction was done using the Benjamini-Hochberg method on probes which passed the significance threshold individually. When assessing contribution of those DMPs to DMRs, P-values represent the percent of permuted regions with more extreme methylation than the null distribution; the family-wise error rate (FWER) values represent the proportion of permutations with at least one region with more extreme methylation than the null distribution^60,68^.

When interpreting analyses of direct paths present in the network structures, Wilcoxon and Fisher’s exact tests were used. Wilcoxon tests were performed to assess the significance of differences between distributions; Fisher’s exact tests were used to assess differences in contingency tables.

We used a Bayesian approach to model the differences in entropy between hypergraph structures in pathways identified as being significantly enriched between KS1 and controls. We iterated 100 hypergraphs each, from genes attributed to the significant pathways and calculated entropy on each of these networks. The distribution of entropy values was resampled 10000 times and the difference between posterior distributions was compared. When the 89% credible interval of the difference between the two distributions did not include 0, the difference was defined as significant. Results were plotted for pathways demonstrated to be significant.

### Gene ontology and epigenetic landscape

Gene ontologies associated with DMPs, DMRs and elements refined from the hypergraph approaches were assessed using WEB-based Gene SeT AnaLysis Toolkit (WebGestalt 2019)^69^. UCSC Genome browser database was used to study the epigenetic landscape (H3K4me1, H3K4me3 and H3K27ac) of selected DMRs^70^ in human blood cells (B cell, CD133HSC, Neutrophils) (Bernstein Lab, Broad Institute, ENCODE consortium).

### Hypergraph analysis

Correlation matrices were generated using R 3.4.1 to investigate the relationships between DMPs, which distinguish KS1 from controls, and all other CpGs that are not differentially methylated. In doing so, a bipartite network is generated which describes pairwise associations between DMPs and all remaining CpGs. In quantifying correlations shared between DMPs, hyperedges in the hypergraph^8–10^ are defined, identifying causally linked groups of DMPs whose methylation patterns are co-ordinated with thousands of other CpGs. Due to the highly multivariate nature of the hypergraph models, they are not limited to assessing cis-regulatory associations.

To generate the hypergraphs, correlation matrices were binarized using an R cut-off equal to the standard deviation of the absolute correlation values. The resulting matrix *M* describes the presence of associations in a binary manner and is the bipartite graph which represents the incidence matrix of the hypergraph. This matrix was then multiplied by the transpose of itself (*M*^t^) to give the final matrix *M×M^t^* whose values describe the number of correlations any pair of DMPs share across the methylome; this represents the adjacency matrix of the hypergraph. By hierarchical clustering of the hypergraph adjacency matrix, a central cluster can be refined as that with the highest number of correlations shared between DMPs. The specific CpGs with which pairs of DMPs share correlations can also be identified from the incidence matrix, to implicate a wider element of the methylome as causally associated with KS1. To do this, we refined the incidence matrix of the hypergraph to a set CpGs which are associated with a large proportion of the target DMPs (>40%).

### Non-negative matrix factorisation (NMF)

NMF clusters the correlation matrix of DMP co-ordination based on the contribution of each DMP to an underlying linear model^71^. This approach can be used to measure co-ordination between DMPs and represents a computationally intensive validation of the hypergraph approach presented in this study.

A correlation matrix was generated between the target DMPs and the remaining CpGs, limited to the 100,000 CpGs with the smallest P-values due to computational constraints. NMF was performed and the resulting matrix was clustered to refine sets of DMPs with similar relationships to one another when compared to the remaining CpGs. We investigated the overlap between clusters generated by NMF and those generated by hypergraph approaches.

### Entropy

Entropy is a mathematical property quantifying organisation. In information theory, Shannon entropy can be used to describe the range of expected outcomes, expressed as a probability distribution of the outcomes. In gene regulatory networks, connections between vertices represent the existence of possible associations between genes; hypergraphs allow us to infer causality in those associations and entropy provides a metric by which we can describe the resulting probability distribution. Network entropy has been demonstrated as informative in defining cellular differentiation potential^33^ and in inferring causal relationships between network elements^56^. Specifically, high entropy describes more uniform distributions and has been associated with less differentiated cellular states^33^.

Shannon entropy was calculated within the central cluster of the adjacency matrix of each hypergraph. The values were compared to those calculated from hypergraphs of randomly selected CpGs, to assess whether co-ordination of DMPs was different to the rest of the methylome. This was iterated 1000 times for both KS1 and controls.

### Network silencing

To assess the underlying map of DMP co-ordination, analyses were performed to refine the direct elements from the hypergraph structure. This approach uses matrix transformation to discriminate direct from indirect associations and potentially refines functional or causal relationships^3^ between network elements. We utilized the method introduced by Barzel and Barabási **(Supplementary Formula 1)**, to silence indirect connections in the hypergraph structure. We then converted the remaining values into Z-scores to assess the distribution of direct network connections in KS1 and controls.

Having assessed differences between the directly connected elements in KS1 and controls, we identified elements which were present uniquely in each group. These unique elements were assessed for associations with known biology via gene ontology analyses. Finally, we assessed the proximity of DMPs defined as directly associated to determine the distribution of cis- vs trans-regulatory associations. We defined cis-associations between DMPs as those where the elements were less than 1Mb apart; elements more distant than this were labelled trans-associations.

## ACKNOWLEDGMENTS AND FUNDING SOURCES

S.C., S.J.K., A.S. and S.B. acknowledge the support of Newlife Charity (grant number 16– 17/10). This work was also supported by GOSH charity (grant number V4621), NIHR Manchester Biomedical Research Centre (NIHR203308 and the MRC Epigenomics of Rare Diseases Node (MR/Y008170/1) and ESPE fellowship (T.G.).

## AUTHOR CONTRIBUTIONS

A.S, S.B and S.J.K conceived and managed work presented. Project carried out by S.C, T.G, E.M. Preliminary analyses carried out by B.W. Manuscript written by S.C and T.G, supported and edited by the other authors. All authors reviewed the final manuscript.

## Notes

**CONFLICT OF INTEREST NOTIFICATION** The authors declare no conflict of interest. The funders had no role in study design, data collection and analysis, decision to publish, or preparation of the manuscript.

### Competing Interest Statement

The authors have declared no competing interest.

