## Supplementary Figures for "Analysis of higher order interactions quantifies co-ordination in the epigenome and reveals novel biological relationships in Kabuki syndrome"

### Supplementary Figure S1. Epigenetic landscape of regions corresponding to the DMRs identified in KS1

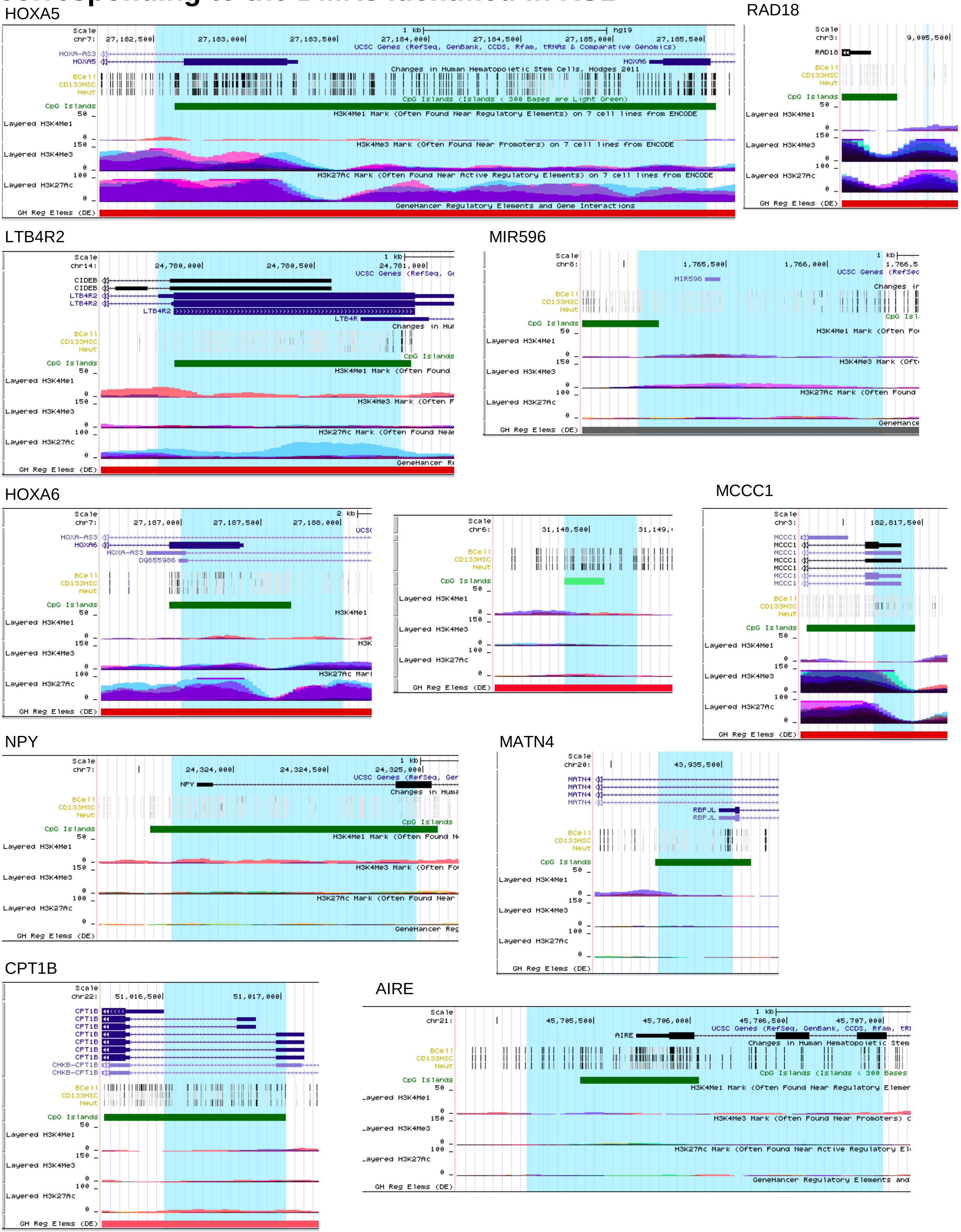

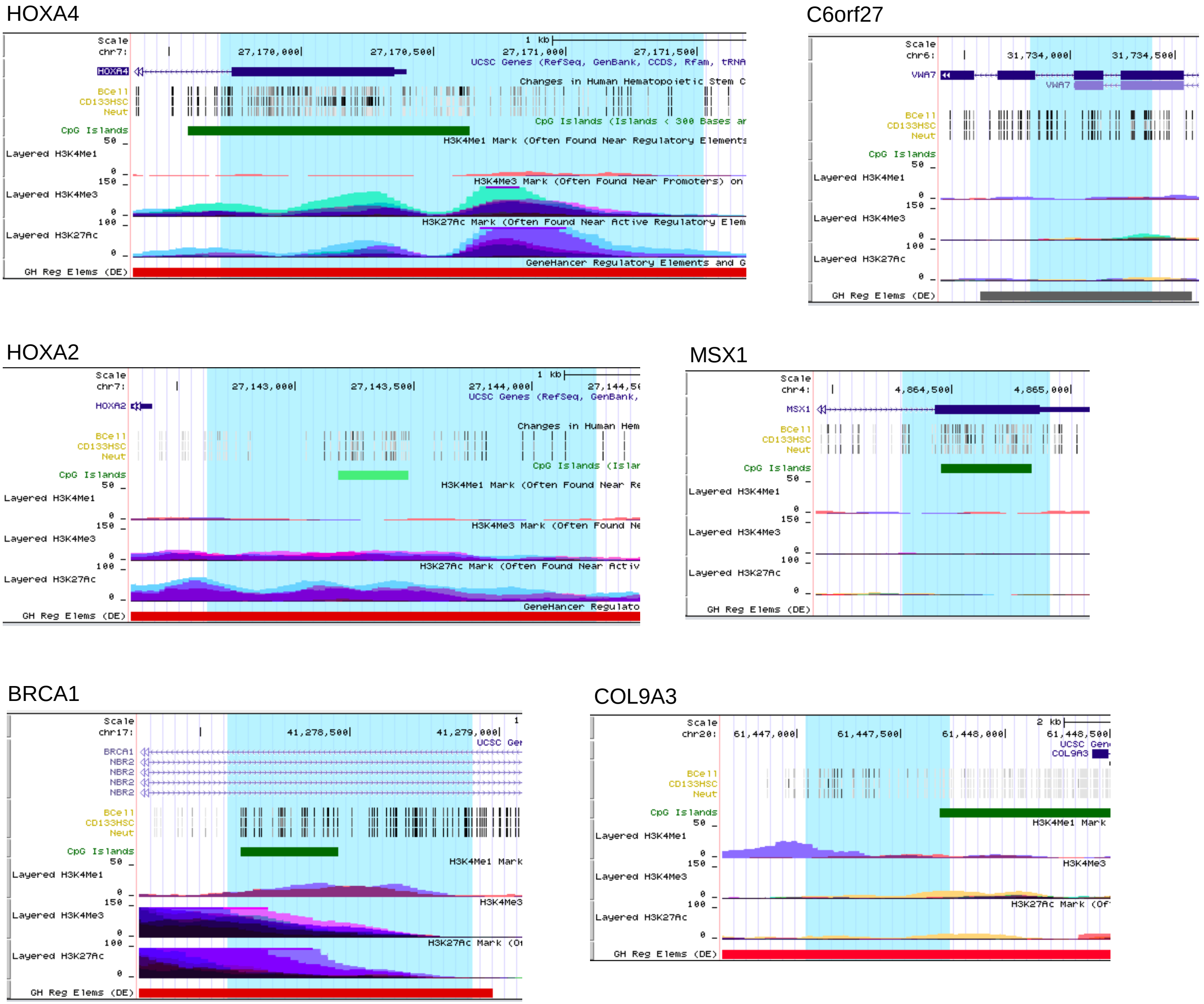


**Supplementary figure 1. Epigenetic landscape of regions corresponding to the DMRs identified in KS1**

Epigenetic landscape (H3K4me1, H3K4me3 and H3K27ac) on genes corresponding to the DMRs identified in KS1 (UCSC Genome Browser).

List of tracks: Chromosome, gene name, methylation level in human blood cells (B cell, CD133HSC, Neutrophils), CpG island, H3K4me1,H3K4me3 and H3K27ac in human blood cells, regulatory element.

**Supplementary table 1. Epigenetic landscape of regions corresponding to the DMRs identified in KS1**

| hypo | H3K4me1 | H3K4me3 | H3K27ac |
|---|---|---|---|
| HOXA4 | - | + | + |
| HOXA2 | - | + | + |
| MSX1 | - | - | - |
| BRCA1 | + | + | + |
| C6orf27 | - | - | - |
| COL9A3 | + | - | - |
| **hyper** | **H3K4me1** | **H3K4me3** | **H3K27ac** |
| HOXA5 | - | + | + |
| LTB4R2 | - | - | + |
| MIR596 | - | - | - |
| HOXA6 | - | + | + |
| NPY | - | - | - |
| MATN4 | - | - | - |
| CPT1B | - | - | - |
| AIRE | - | - | - |
| RAD18 | - | + | + |
| MCCC1 | - | + | + |
| DMR7 | - | - | - |

### Supplementary Figure S2. Analysis of co-ordinated interactions between DMPs in KS1 using NMF

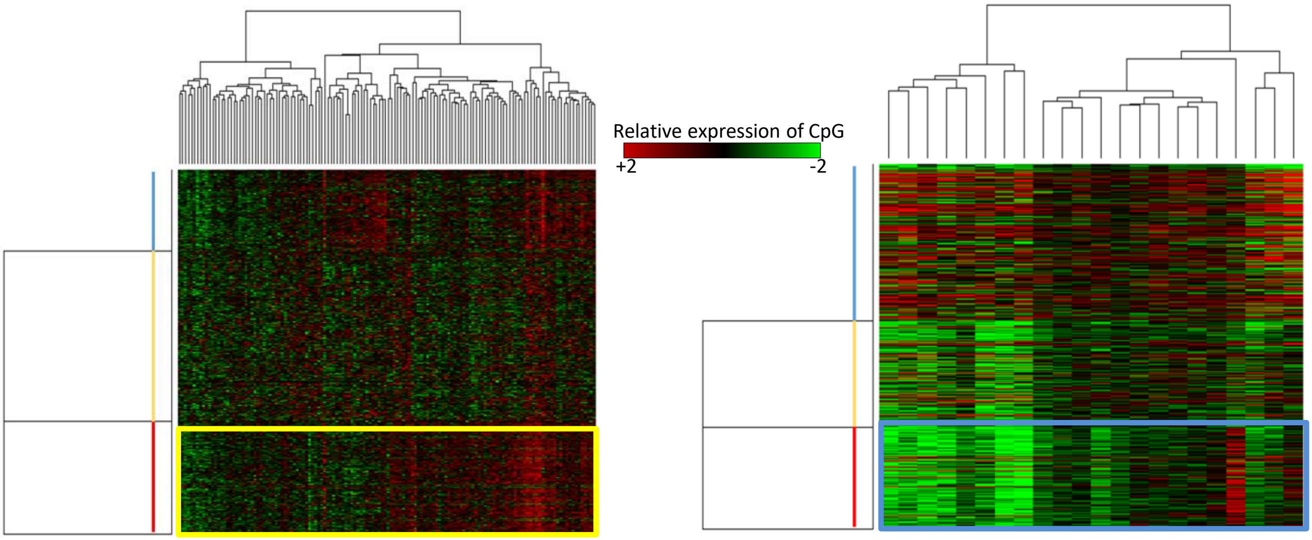


**Supplementary Figure S2. Analysis of co-ordinated interactions between DMPs in KS1 using NMF.** Clusters of DMPs identified by non-negative matrix factorisation (NMF) [blue, yellow, red]. The red cluster demonstrated >90% similarity to the hypergraph data cluster (yellow and blue boxes in Figure 4).

### Supplementary figure S3. GO analysis on central cluster DMPs in controls and KS1

**A** **Control**

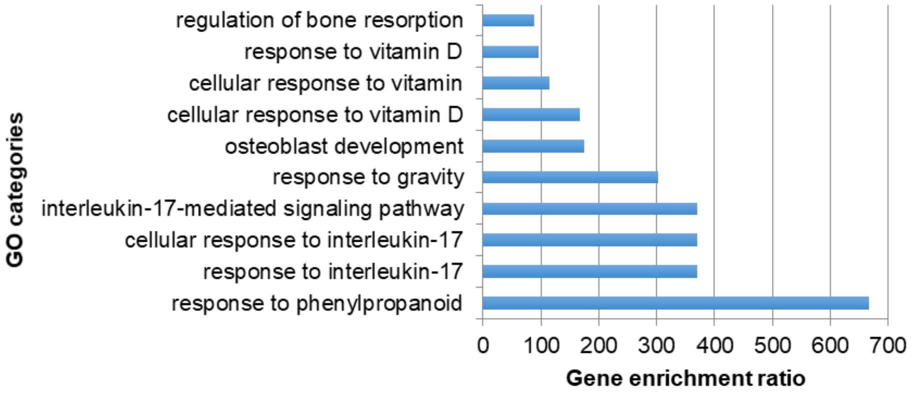


**B** **KS1**

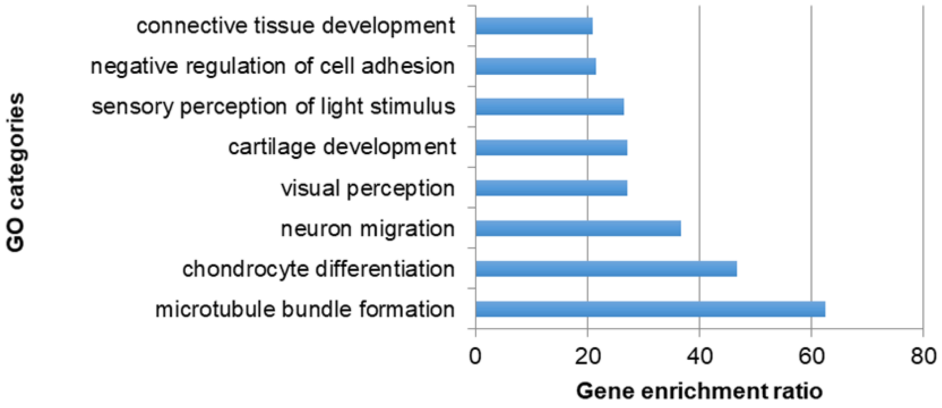


**Supplementary Figure S3. GO analysis on central cluster DMPs in controls and KS1**
List of gene ontology categories performed on central cluster DMPs in control (**A**) and KS1 (**B**) identified by WebGestalt (FDR >0.05, P< 0.05). In the control 7 of 1036 IDs mapped to unique entrezgene IDs while in the KS1 3 out of 986 IDs mapped to unique entrezgene IDs.

### Supplementary Figure S4. Gene ontology associated entropy differences in the hypergraph between KS1 and controls

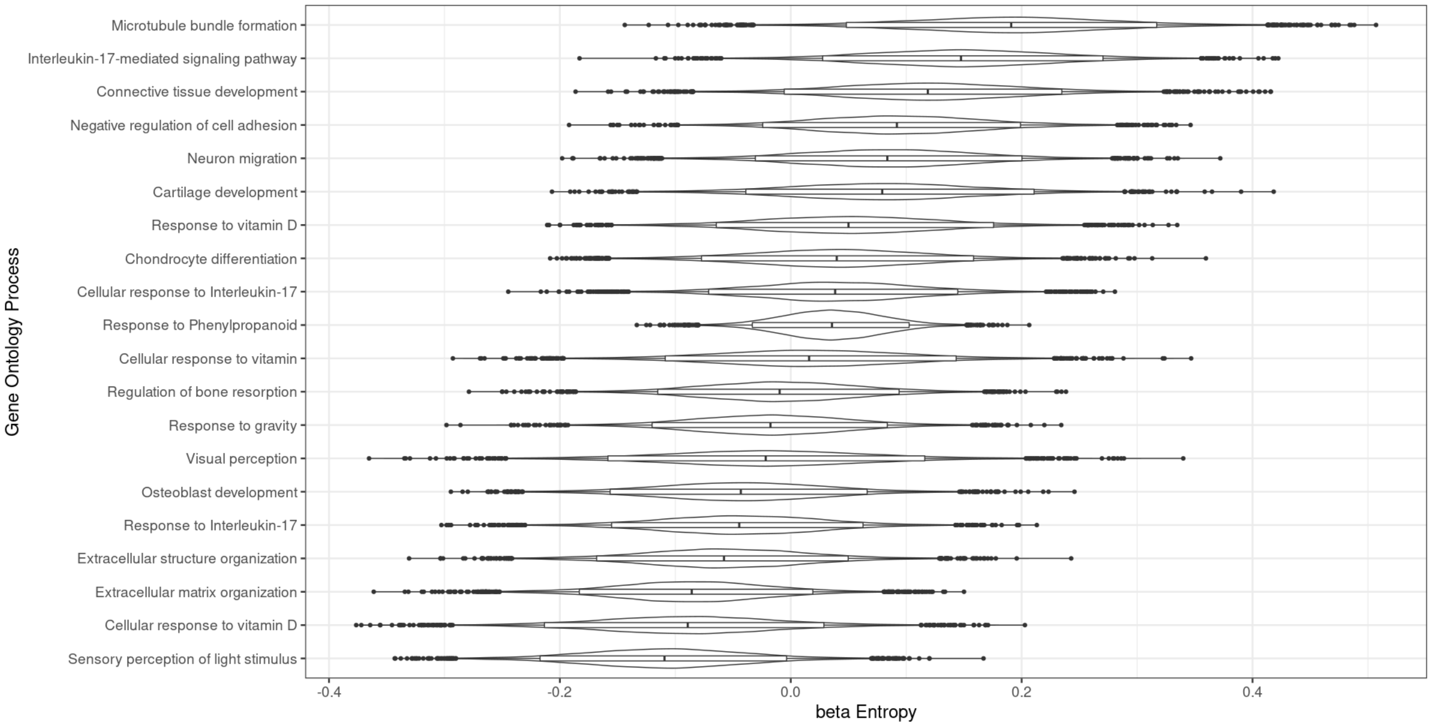


**Supplementary Figure S4. Difference in entropy between processes identified as significantly associated with hypergraph central cluster DMPs.** Bayesian MCMC simulation of distributions of hypergraph entropy for a range of processes identified as significantly associated with DMPs from the KS1 and control hypergraph central clusters. hypergraphs were iterated using DMPs in genes associated with each biological process and distributions were resampled. Differences between KS1 and control were modelled using a Bayesian MCMC approach and differences were determined to be significant in those where the 89% credible interval of the difference between groups (beta) did not encompass 0. Those processes were selected for figure 5C.

### Supplementary Figure S5. Gene ontology associated direct paths in the hypergraphs of KS1 and controls

**A) Controls**

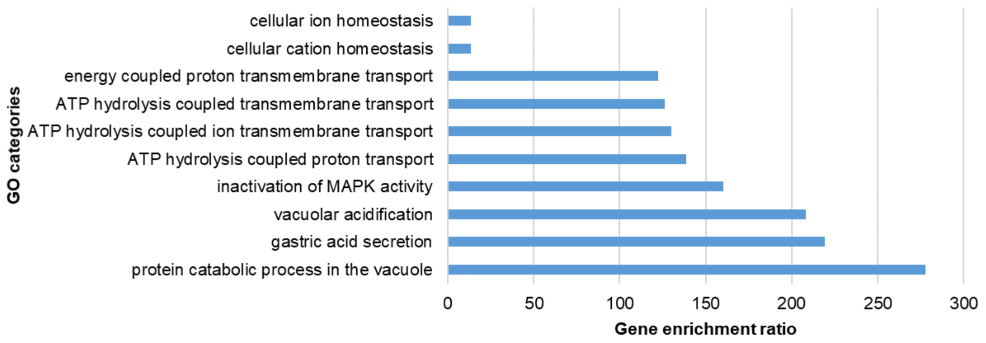


**B) KS1**

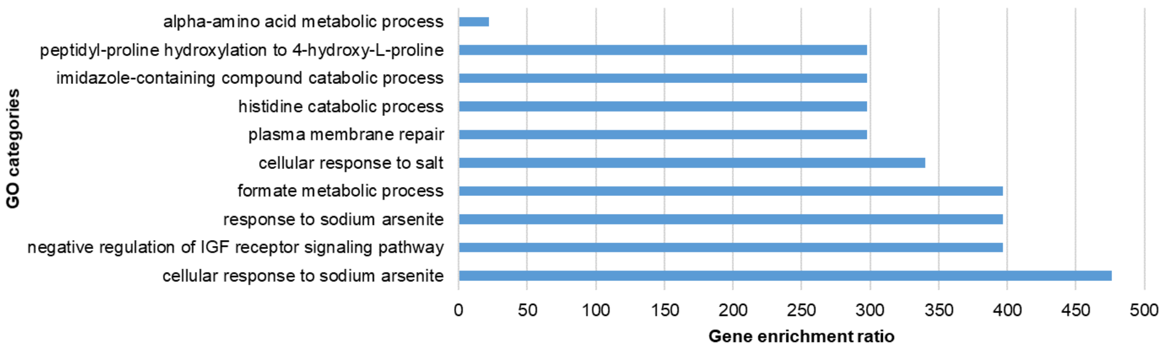


**Supplementary Figure S5. Gene ontology associated direct paths in the hypergraphs of KS1 and controls**

List of gene ontology categories performed on  in control (**A**) and KS1 (**B**) identified by WebGestalt software (FDR>0.05, P<0.05). In the control 5 out of 14 IDs mapped to unique entrezgene IDs while in the KS1 7 out of 19 IDs mapped to unique entrezgene IDs.

**Supplementary Formula SF1. Network link prediction formula to silence indirect associations in the hypergraph structure.**

$$S = (G - I + \mathcal{D}((G - I)G))G^{-1}$$

S = Silenced matrix
G = hypergraph adjacency matrix
I = Identity matrix
$\mathcal{D}$(M) = a function which sets the off-diagonal terms of M to zero
